# Isolation and manipulation of meiotic spindles from mouse oocytes reveals migration regulated by pulling force during asymmetric division

**DOI:** 10.1101/2024.12.06.627260

**Authors:** Ning Liu, Ryo Kawamura, Wenan Qiang, Daniela Londono, Ahmed Balboula, John F Marko, Huanyu Qiao

## Abstract

Spindles are essential for accurate chromosome segregation in all eukaryotic cells. This study presents a novel approach for isolating fresh mammalian spindles from mouse oocytes, establishing it as a valuable *in vitro* model system for a wide range of possible studies. Our method enables the investigation of the physical properties and migration force of meiotic spindles in oocytes. We found that the spindle length decreases upon isolation from the oocyte. Combining this observation with direct measurements of spindle mechanics, we examined the forces governing spindle migration during oocyte asymmetric division. Our findings suggest that spindle migration is regulated by a pulling force and a net tensile force of approximately 680 pN is applied to the spindle *in vivo* during the migration process. This method, unveiling insights into spindle dynamics, holds promise as a robust model for future investigations into spindle formation and chromosome separation. We also found that the same approach could not isolate spindles from somatic cells, indicative of mammalian oocytes having a unique spindle organization amenable to isolation.

**Significance Statement:** Spindles are essential for accurate chromosome segregation in all eukaryotic cells, yet studying their mechanical properties in vivo remains challenging. Here, we present a novel method for isolating intact, functional spindles from live mouse oocytes, establishing a powerful in vitro model for dissecting spindle mechanics. Using this system, we reveal that spindle length shortens upon isolation and quantify a net tensile force of approximately 680 pN applied to the spindle during migration, implicating a pulling mechanism in asymmetric division. Notably, this isolation approach does not succeed in somatic cells, highlighting a unique organization of the mammalian oocyte spindle. This platform opens new avenues for understanding spindle dynamics, force generation, and chromosome segregation in meiosis.

## Introduction

The spindle serves as an essential structure for chromosome separation and cell division, yet many questions remain regarding its components and structure (1). Many approaches have been developed to investigate the spindle, with one of the most powerful methods being the utilization of *Xenopus* egg extracts (2, 3). This approach involves extracting cytoplasm from *Xenopus* eggs, supplemented with a sperm nucleus. Upon activation, this combination is able to form a spindle structure (4, 5). This system has been used to identify many spindle components and understand spindle formation processes (6, 7), and can be synergistically employed with other techniques to study spindle characteristics (8, 9). However, *in vitro* egg-extract spindles may differ from *in vivo* spindles as the former are assembled in a cell-free system. Additionally, this method is exclusive to amphibians due to the necessity of collecting numerous oocytes to obtain sufficient cytoplasm volume, which is much more challenging to obtain from mammalian cells. New methods applicable to investigate mammalian spindles are needed.

In the realm of spindle research, most studies have been conducted in cells rather than *in vitro*, which makes it problematic to accurately understand spindle physical properties within the intricate cytoplasmic environment. Some previous attempts involved isolating spindles from somatic cells after stabilizing microtubules with taxol (10). This isolation process requires physically and chemically stressful treatments, which may alter spindle components and structure, leading to differences between the *in vitro* and *in vivo* spindles. In this study, we introduce a novel method that integrates oocyte *in vitro* culture with micromanipulation to efficiently isolate fresh, intact spindles from mammalian oocytes in a quick and straightforward manner, avoiding use of microtubule-altering drugs. We use this approach to provide direct measurement of oocyte spindle mechanics, which leads us to infer that the spindle is subjected to appreciable stress *in vivo*. This method also offers the advantage of obtaining fresh spindles at different substages based on an oocyte culture timeline.

## Results

The oocyte spindle is subjected to a pulling force during spindle migration *in vivo* Previously studies have shown that F-actin (11, 12) and metaphase cytoplasmic microtubule organization center (mcMTOC) locate on opposite sides of the oocyte spindle, generating tension that stretches the spindle apparatus (13). Unlike the symmetric division observed in somatic cells and spermatocytes, oocytes undergo asymmetric division, requiring the spindle to migrate from the cell center to the cortex. This migration has sparked debate over whether it is driven primarily by pulling forces (11, 12) or pushing forces (14–16).

To address this, we disrupted the forces exerted on both sides of the spindle and observed changes in spindle morphology in mouse oocytes. Laser ablation of the mcMTOC led to a significant reduction in spindle length, which indicates that the mcMTOC contributes to spindle stretching *in vivo* (pre-ablation: 30.6 ± 0.5 μm versus post-ablation: 25.7 ± 0.7 μm, *P* < 0.0001, t-test) (Fig. 1 A and B). Similarly, disrupting F-actin with cytochalasin D, an actin polymerization inhibitor (17), resulting in a significant reduction in spindle length after just 2 minutes of treatment (pre-treatment: 33.7 ± 0.8 μm versus post-treatment: 32.0 ± 0.9 μm, *P* < 0.01, paired t-test) (Fig. 1 C). Collectively, these results suggest that spindle migration during oocyte meiosis is primarily governed by pulling forces that generate tension along the spindle axis. If a pushing force were driving spindle migration, the spindle should be compressed, yielding a shorter length *in vivo* than after isolation when cellular constraints are released.

**Figure 1.**
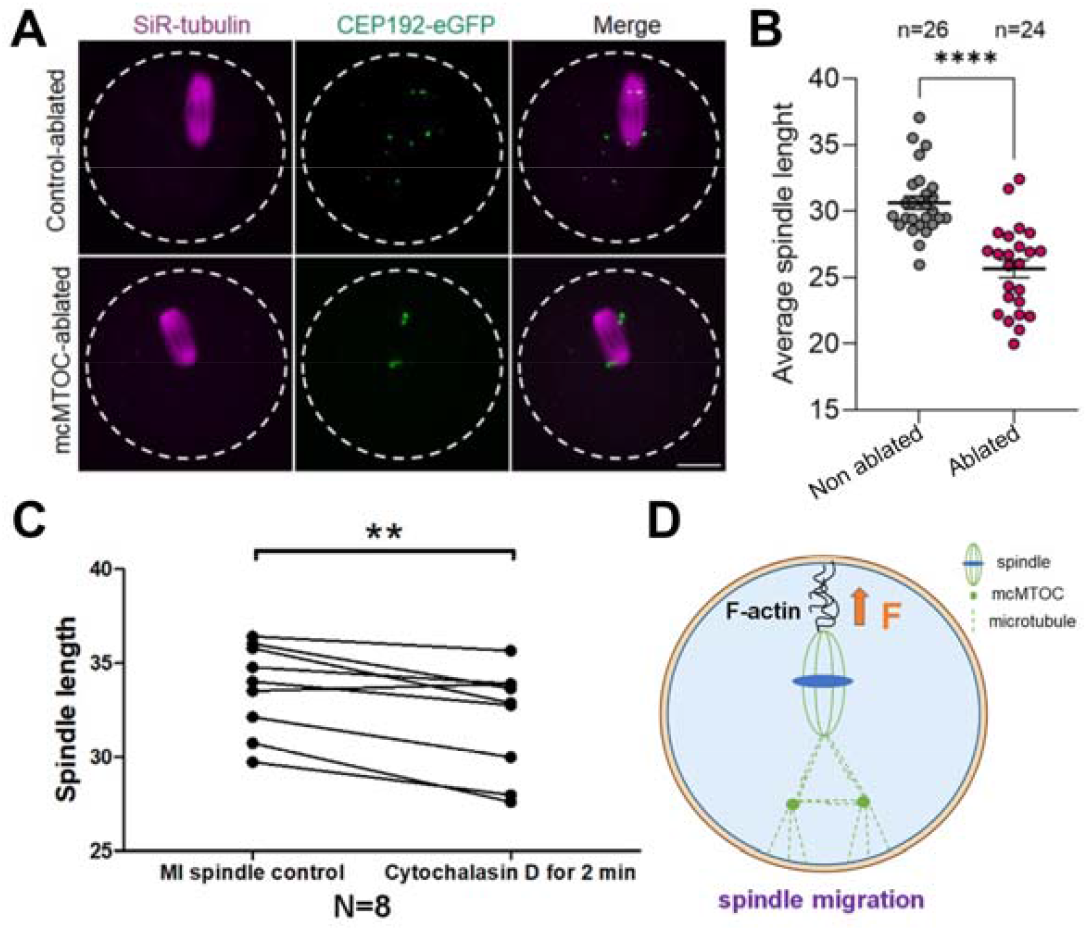
Spindle migration is driven by pulling forces rather than pushing forces. (A) Representative images from mcMTOC laser ablation experiments. (B) Spindle length decreases *in vivo* following mcMTOC ablation. (C) Spindle length decreases after two minutes of cytochalasin D treatment *in vivo*. (D) The MI spindle withstands stretching forces inside oocytes, and its migration is driven by pulling forces. The oocyte spindle migrates from the center to the cortex, likely driven by pulling forces mediated by F-actin, while the opposite side is anchored by the mcMTOC.

### Efficient *in situ* isolation of meiotic spindles using micromanipulation

Oocyte spindles exhibit phase-separation property (18), which enable their structural integrity to be maintained during isolation. This unique characteristic allows the spindle to be isolated intact, providing an opportunity to disrupt the pulling forces normally exerted from both ends. Such isolation offers an independent approach to test whether pulling, rather than pushing, drives spindle migration. We hypothesize that disrupting these forces via spindle isolation would lead to a reduction in spindle length, similar to the effects observed following mcMTOC ablation and cytochalasin D treatment.

Previously, we successfully used micromanipulation to isolate prophase I and metaphase chromosomes (19, 20). Using a similar approach, we isolated intact Metaphase I (MI) spindle from oocytes after 6 hours of culture (Fig. 2 A). The process began with removal of the zona pellucida (Fig. S1 A arrow) using acidic Tyrode’s solution, followed by recovery in PBS (Fig. S1 A). The MI spindle appeared as a barrel-shaped structure containing dark, short threads – indicative of chromosomes – at its center (Fig. 2 B left). Next, we gently microsprayed 0.05% Triton X-100 solution to generate a hole in the oocyte membrane. The MI spindle spontaneously flowed out of the oocyte. Using phase contrast imaging, the microtubule fibers and typical meiotic chromosomes were visible (Fig. 2 B right). This approach enables spindle manipulation under phase contrast. In addition, the detailed structure of isolated spindles could be observed from any angle to examine the chromosome and microtubule organization (Fig. 2 C). To validate the isolated spindle identity, we microsprayed α-tubulin antibody to stain the microtubules/spindle and used Hoechst to stain the DNA/chromosomes. After staining, many parallel microtubules within the spindle with well-aligned chromosomes/DNA in the middle were observed, confirming the successful isolation of the spindle (Fig. 2 D).

**Figure 2.**
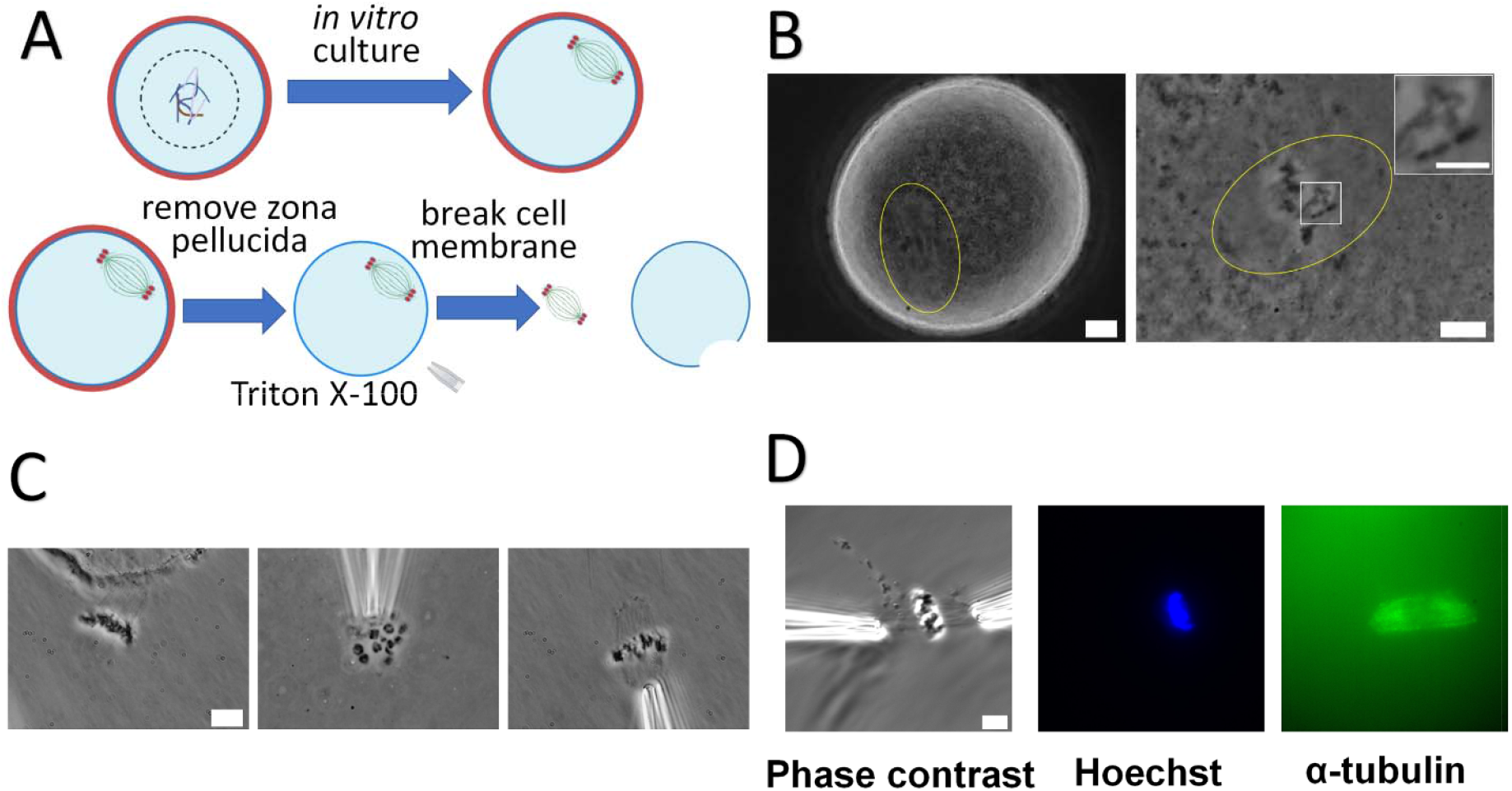
Spindle isolation and confirmation. (A) Cartoon of oocyte spindle isolation (B) Spindle isolation from oocytes whose zona pellucida was removed. Left panel: An oocyte immersed in PBS after zona pellucida removal. The spindle outline (circled with yellow) can be observed under the phase contrast microscopy. Scale bar=10 μm. Right panel: A MI spindle after isolation, showcasing the spindle structure (circled with yellow) and meiotic chromosomes with chiasmata (circled with white). Scale bar=10 μm. Right panel upper right: An enlarged image of the meiotic chromosomes with chiasmata. Scale bar=5 μm. (C) MI spindle isolation. Left panel: spindle comes out of the oocyte after the cell membrane is broken. Middle panel: vertical review of the spindle (top view). Right panel: horizontal view of the spindle (side view). Scale bar=10 μm (D) Spindle confirmation. Left panel: phase contrast image. Middle panel: Hoechst staining for DNA (Blue). Right panel: α-tubulin staining for microtubules (Green). Scale bar=10 μm.

We found that the choice of extraction buffer is important to preservation of spindle structure. As an example, that illustrates this, we also tested PEM (general tubulin buffer) solution which is often used to stabilize microtubules *in vitro* (21). We found that this type of low-salt PEM solution is highly toxic to oocytes and alters their morphology within a few seconds (Fig. S2 A). Although osmotically balanced PEM solution (classical PEM plus about 150 mM univalent salt) does not rapidly disrupt oocyte morphology, isolated spindles in osmotically balanced PEM disassemble much more rapidly than spindles extracted into PBS (Fig. S2 B and S3 A). We isolated spindles from oocytes and monitored their morphological change over time in PBS (video 1). We observed the spindles to be stable, exhibiting minimal morphological changes for 30 minutes following isolation (Fig. S3 A). Overall, PBS maintains the structure of isolated spindles, and in this regard is superior to PEM-based buffers.

### Isolation of meiotic spindles at different stages

The timeline of the oocyte *in vitro* maturation has been heavily studied and is illustrated in Fig. 3 A. In addition to MI oocyte isolation, our method is also applicable for the isolation of metaphase II (MII) spindles after 14 hours of oocyte culture (Fig. 3 B). The high stability of isolated spindles opens up new avenues for studying spindles in both meiosis I and meiosis II. Compared to MI spindles, MII spindles did not easily flow out of MII oocytes after oocyte-membrane rupture and exhibited a stronger mechanical attachment to the cell membrane, making their isolation more challenging (22).

**Figure 3.**
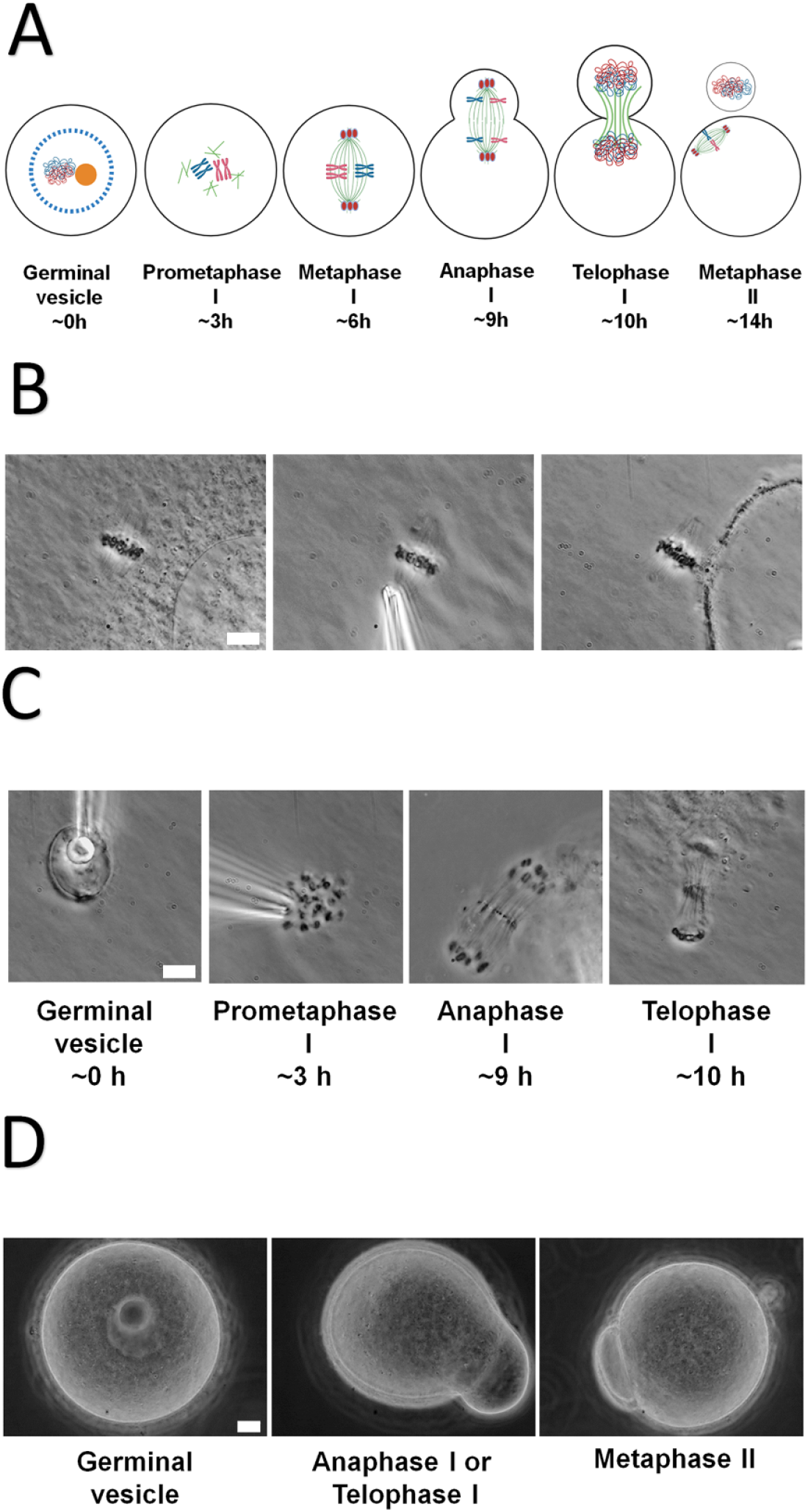
Isolation of spindles at different meiotic stages. (A) Spindle formation process and timeline during oocyte meiosis (B) MII spindle isolation. Left panel: MII spindle after isolation from oocyte. Middle panel: MII spindle held by glass pipette. Right panel: MII spindle tightly connected to the cell membrane. Scale bar=10 μm (C) Spindle isolation at other stages. At around 3h, the spindle ball with condensed chromosomes can be isolated. At around 9h, the anaphase spindle, with separated homologous chromosomes moving toward different poles, can be isolated. At around 10, the telophase spindle, with an hourglass shape and decondensed chromosomes, can be isolated. Scale bar=10 μm (D) Oocyte features can be used to distinguish different stages. For oocyte at germinal vesicle stage, large nucleoli are visible under phase contrast microscopy. For the oocyte at anaphase I or telophase I stage, it is still connected with a small bud. For oocyte at MII stage, the first polar body is completely extruded from the oocyte. Scale bar=10 μm.

Our method could be extended to study spindles at other substages of meiosis simply by adjusting the *in vitro* culture duration according to the desired substage along the timeline. For example, it is feasible to isolate prometaphase pre-spindle bundles, anaphase I and telophase I spindles (Fig. 3 C). Spindle-stage determination can be facilitated by oocyte shapes and characteristics during isolation (Fig. 2 B and 3 D). Oocytes at the diplotene/germinal vesicle (GV) stage exhibit prominent nucleoli (Fig. 3 D left). At metaphase I stage, the nucleolus is absent, and the spindle is visible under phase contrast microscopy (Fig. 2 B). Anaphase I is characterized by a small bud emerging from the oocyte (Fig. 3 D middle). At metaphase II stage, a fully separated polar body is evident (Fig. 3 D right).

The rapid and simple isolation method yields fresh spindles without the need for more physical or chemical treatments (e.g., osmotic stress lysis, centrifugation), preserving their original state as much as possible. Most importantly, our approach can repetitively produce well-regulated spindles in a stable manner, as the spindles are formed within live oocytes. In our experiments, we successfully isolated spindles from 95 out of 106 (∼90%) metaphase I oocytes using this method. Among these failed isolations, four resulted from the isolation failure, where the spindle formed (Fig. S1 B left circled with yellow) but could not be isolated for some unknown reasons. The remaining seven were due to spindle-formation delay, where the spindles had not fully developed at the time of isolation (Fig. S1 B right). Overall, this method provides a consistently reliable way to obtain high-quality, fresh spindles from mouse oocytes. Moreover, it is readily adaptable for cell biology studies, allowing oocytes to be exposed to different chemicals or drugs during culture, followed by spindle isolation to investigate effects at any meiotic stage.

### Manipulation of isolated meiotic spindles *in vitro*

After spindle isolation, meiotic spindles exhibit remarkable stability, allowing them to persist in PBS for an extended duration. The spindles can be easily manipulated with two micropipettes, either from the side or ends (Fig. 4 A). Our attempts to manipulate the isolated spindles revealed intriguing insights (Fig. 4 D). Stretching spindle on the side resulted in the longitudinal disintegration of the spindle, revealing individual microtubule bundles and confirming the spindle’s composition of inter-connected microtubules (23) (Fig. 4 B arrow). Stretching spindles on the ends led to the separation of the two half-spindles, albeit still connected by chromosomes (Fig. 4 C). With further stretching, the two half-spindles separated from one another, and all chromosomes were subsequently removed from one half spindle. Notably, both half-spindles remained stable and intact, which means the spindle structure is independent of the connected chromosomes. This observation aligns with the concept that the spindle can be viewed as two half-spindles connected by chromosomes (1, 24).

**Figure 4.**
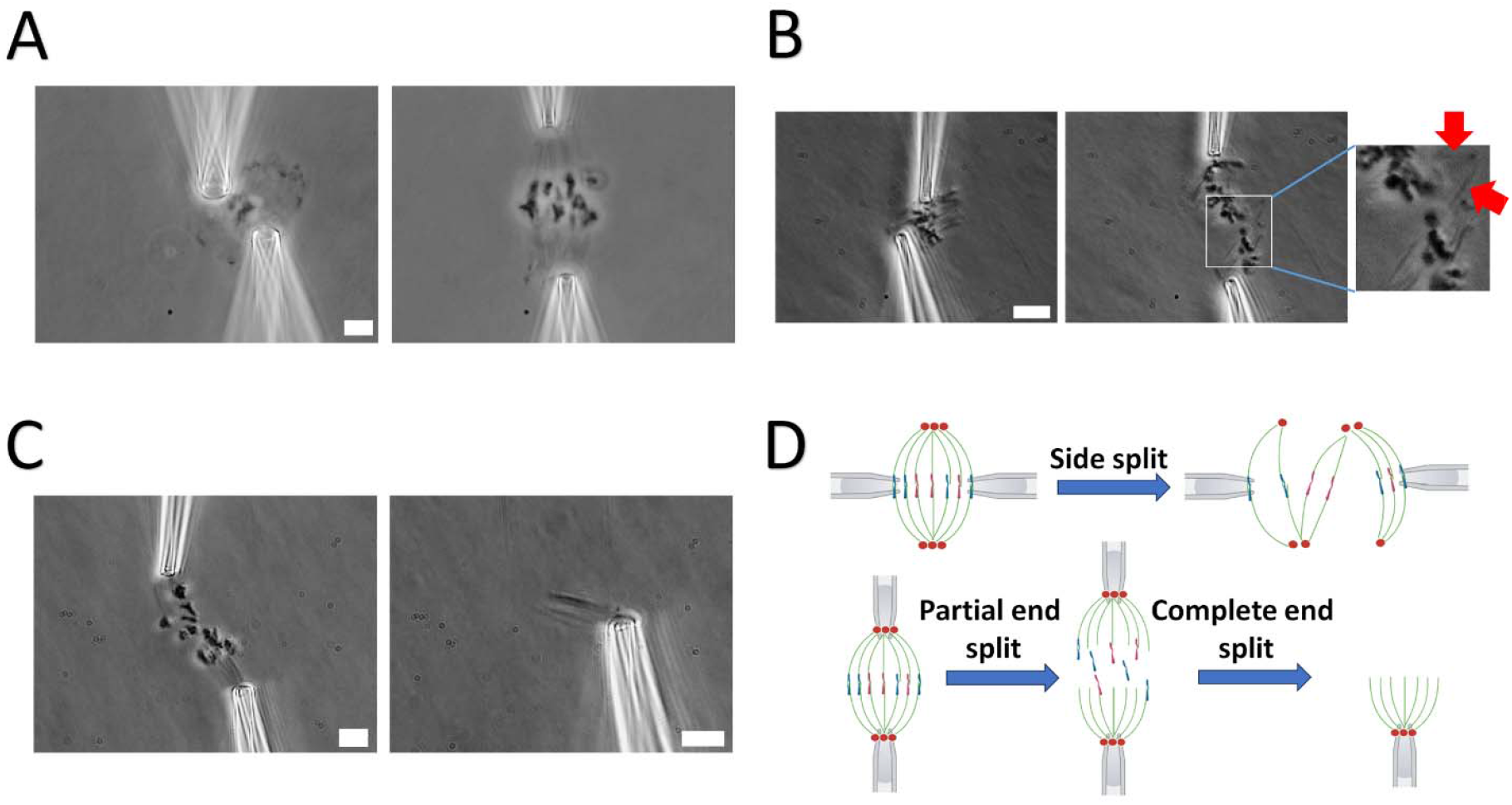
Manipulation of isolated meiotic spindles. (A) Spindles can be grabbed at different angle using two micropipettes. Left panel: spindle is grabbed on the side. Right panel: spindle is grabbed at the ends. Scale bar=10 μm (B) Stretching spindle on the side could split microtubule arrays. Left panel: before stretch. Middle panel: after stretch. Stretching on side causes spindle disintegration. Some clear microtubules still could be seen (enclosed in white box). Right panel: A enlarged image of split microtubules (red arrow). Scale bar=10 μm (C) Stretching spindle on the ends could break it into two half spindles. Left panel: stretching spindle on the ends. Right panel: stretching to remove all chromosomes from one half spindle. Stretching spindle on the ends could split a whole spindle into two half spindles with chromosomes still connecting them. Moving two half spindles far away could result in all chromosomes being removed from one of the two half spindle, while maintaining spindle structure. This indicates that the spindle structure is independent of chromosomes. Scale bar=10 μm (D) A diagram of oocyte spindle manipulation.

### Strong interaction between centrosome and cell cortex hinders spindle isolation in somatic cells

We attempted to apply this oocyte spindle isolation method to somatic cells (>30), including human HeLa cells, mouse embryonic fibroblasts (MEFs), and newt TVI cells at metaphase stage (Fig. 5 A, B and S4) (25). However, after cell-membrane lysis, we did not observe spindles with chromosome bundles to freely flow out, and spindle isolation proved unsuccessful. This phenomenon suggests that spindles in somatic cells and oocytes are different. We think that the difference of spindle organization between these two systems may be the cause. The two most likely explanations of this are: 1) mitotic spindles become unstable after cell lysis, and 2) a strong connection between the spindle and other cellular components prevents isolation of mitotic spindles.

**Figure 5.**
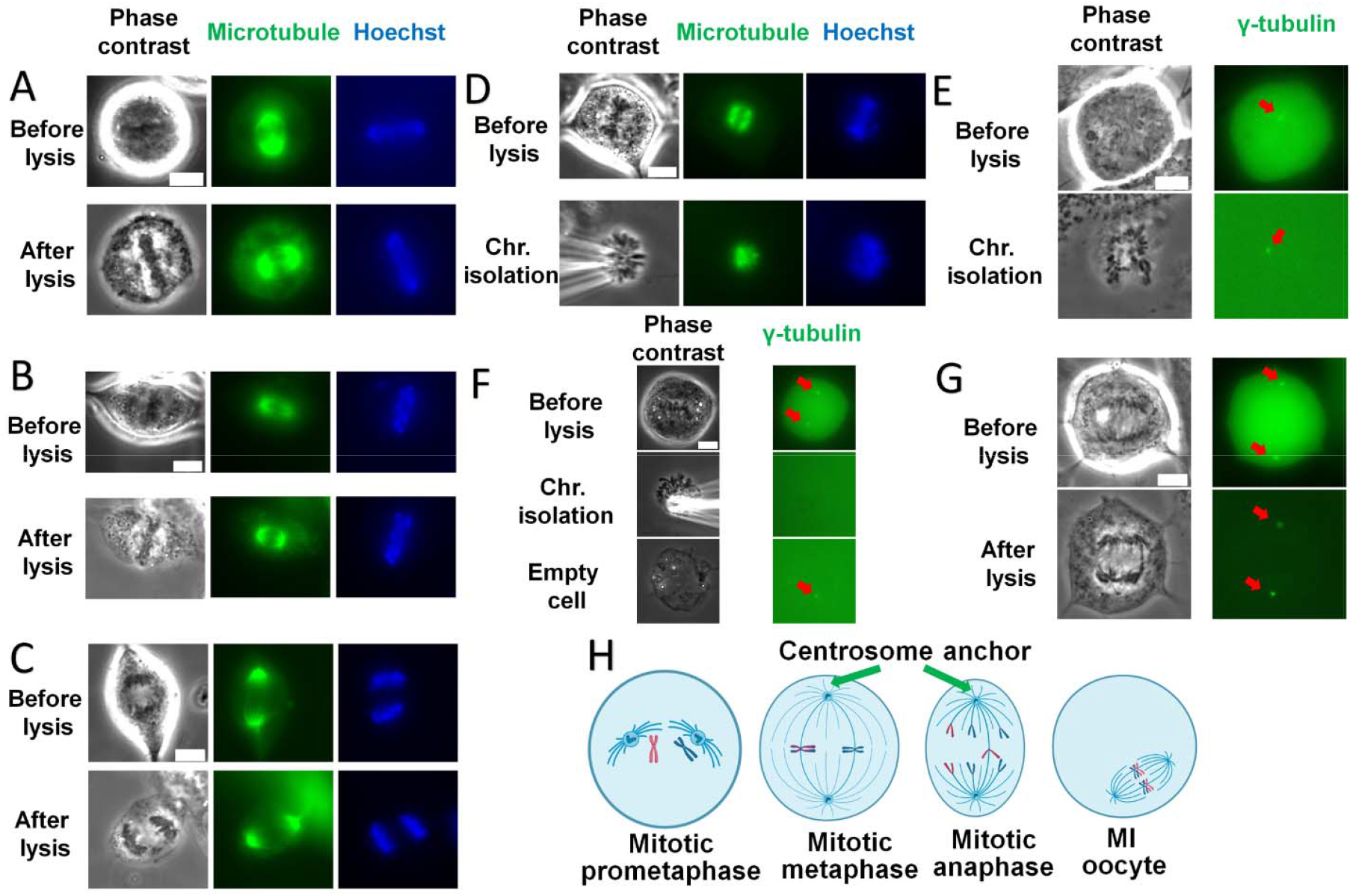
Strong interaction between centrosome and cell cortex. (A-C) Cell images before and after cell lysis. Microtubules are stained in green, and DNA/chromosomes are stained in blue. Metaphase Hela cells (A); Metaphase MEF cells (B); Anaphase MEF (C). Scale bar =10 μm. (D) Prometaphase MEF cells with microtubule staining in green and DNA/chromosome Hoechst staining in blue. The top row shows the intact MEF cells before cell lysis; the bottom row demonstrates that prometaphase spindle with chromosomes can be easily isolated. Scale bar =10 μm. (E) Prometaphase centrosomes highlighted with γ-tubulin staining are isolated with spindle. Prometaphase U2OS centrosome marked by γ-tubulin in green. Left panel: phase contrast. Right panel: γ-tubulin. Scale bar =10 μm. (F) Metaphase chromosome isolation without accompanying centrosomes. Metaphase U2OS cells with centrosome (γ-tubulin) staining. Left panel: phase contrast. Right panel: γ-tubulin. Scale bar =10 μm. (G) Anaphase U2OS cells with microtubule staining. Left: phase contrast. Right: γ-tubulin. Scale bar =10 μm. (H) Interaction between the centrosome and cortex in different types of cells. In somatic cell, during prometaphase stage the chromosomes and centrosomes could be easily isolated from the cell because there is no strong interaction between the centrosome and cell cortex. During mitotic metaphase and anaphase, the centrosome has already formed a strong interaction with cell cortex, preventing spindle isolation. In contrast, oocyte spindle lacks centrosome and astral microtubules connected to cell cortex, resulting in no strong interaction between the spindle and cell membrane, making isolation feasible.

To explore the first possibility, we stained microtubules and observed intact spindle structure in Hela cells and MEF cells after membrane lysis, indicating that mitotic spindles remain stable after cell lysis (Fig. 5 A, B and S5). Our focus shifted to the second possibility – strong connections to other cellular structures hinder spindle isolation.

To test the second hypothesis, we utilized U2OS cells with GFP-labeled γ-tubulin to track the centrosomes. As previously described (19, 26), chromosome bundles could be easily isolated at somatic prometaphase, an early stage without tight connection between centrosomes and cell cortex (Fig. 5 D, E); prometaphase centrosomes came out of the lysed cell along with the chromosome bundles rather than staying inside the cell (Fig. 5 E). In contrast, at mitotic metaphase, the spindles from mitotic metaphase cells stayed inside the cell after its lysis. Attempts to physically drag chromosome bundles out result in isolated bundles without visible attached spindle structure (Fig. 5 F). Furthermore, the centrosome remained inside the cell rather than coming out with the chromosome bundles. These results suggest that chromosome isolation disrupts the connection between chromosomes and the centrosomes, rather than the connection between centrosomes and cell cortex. Previous literature aligns with this result: astral microtubules emanating from centrosomes are anchored to somatic cell cortex via NuMA and dynein (27–30) and centrosome-cortex connection become progressively stronger through mitosis (31).

We concluded that our micropipette-micromanipulation approach was unable to isolate intact spindle structures from somatic cells due to the strong interaction between the centrosome and the cell cortex (Fig. 5 H). Attempts to isolate spindles from somatic cells at the anaphase stage were also unsuccessful, providing further confirmation of the impact of this interaction (Fig. 5 C, G). In contrast, oocytes with acentrosomal microtubule-organizing centers (aMTOCs) lack such a strong interaction, enabling spindle isolation throughout meiosis (Fig. 2 B, 3 B, C and 5 H) (32). Lack of centrosome and astral microtubules in oocytes may explain the ease with which an oocyte spindle can freely flow out after cell lysis.

After cell lysis and chromosome bundle isolation, centrosomes persisted for an extended time (Fig. 5 E, F) suggesting that centrosomes may have a stable and solid structure, consistent with previous published results (33).

### Mechanics of the oocyte spindle

To investigate the stiffness of oocyte spindles, we attempted to stretch the spindles using glass micropipettes. Initial attempts revealed instability in the nonspecific-adhesion connection between the spindles and glass micropipettes, resulting in detachment during stretching. To address this problem, we used specially treated glass micropipettes that adhered to protein-based objects via amine chemistry (Fig. 6 A top & Materials and methods). This modification facilitated a robust attachment of the pipettes to the spindle, enabling controlled stretching without detachment. As we stretched the spindle, a noticeable transformation occurred - narrowing and elongation - which indicated that the spindle is elastic and possesses a deformable gel-like structure (Fig. 6 B, C, D and video 2). The reversible return of the spindle to its original status upon force retrieval further supports its elastic properties (Fig. 6 E). However, excessive stretching led to an irreversible change (Fig. 4 C, 6 D). Throughout the stretching process, where force was only applied onto the ends of the spindle, both chromosomes and microtubules underwent elongation, underscoring mechanical connection to each other. This observation is consistent with the fact that chromosome segregation is due to the external force from microtubule (34).

**Figure 6.**
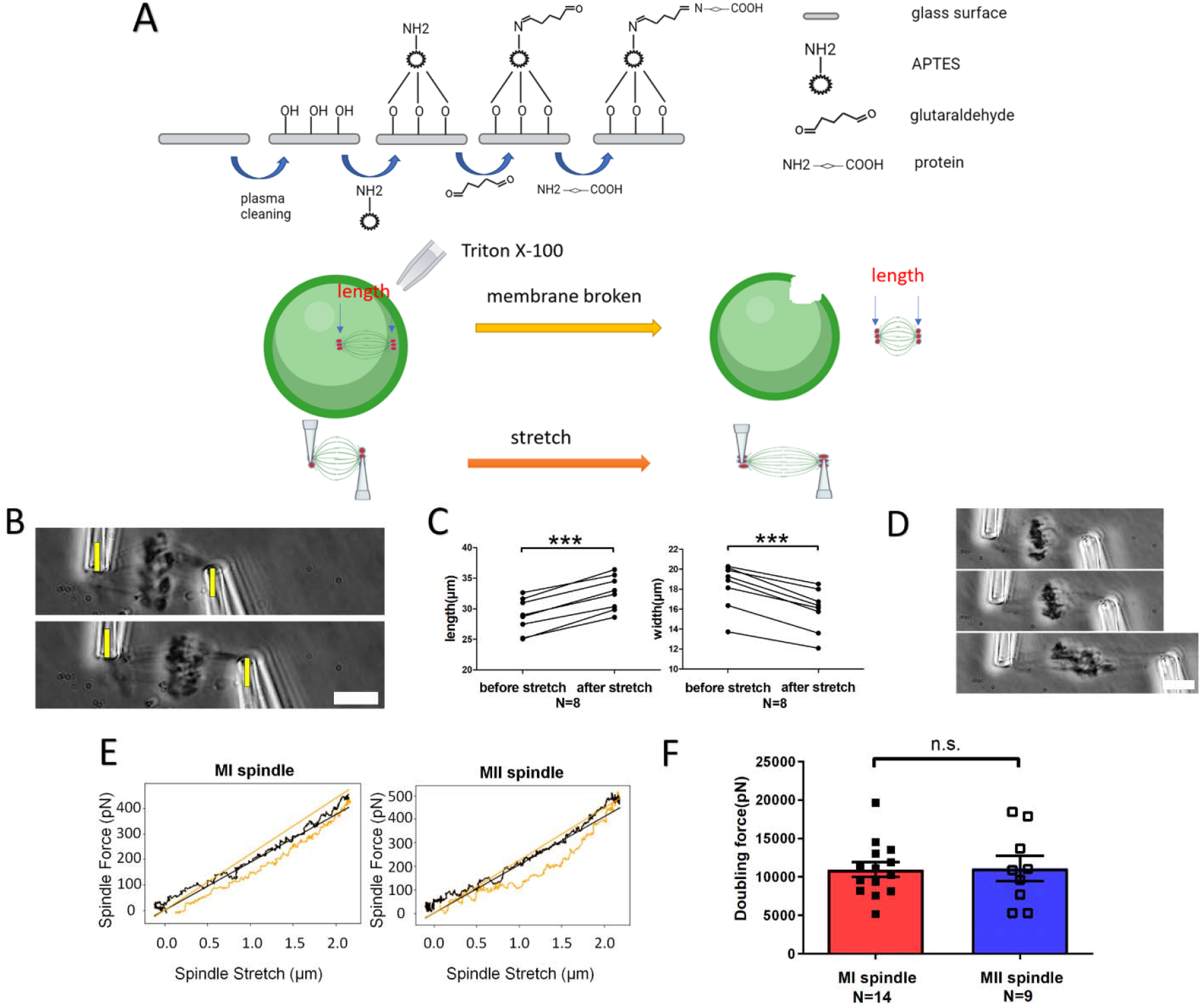
Elasticity measurement of the oocyte spindles. (A) Experiment sketch. Top: pipette preparation. Middle: spindle isolation. Bottom: spindle stiffness measurement. (B) An example of spindle stretching. Top panel: Before stretching. Bottom panel: After stretching. The spindle is elastic, allowing it to be stretched and increasing its length. Scale bar=10 μm (C) Spindle length and width change after stretch. During stretching, the spindle length increases while its width decreases. (D) Excessive spindle stretching. Scale bar=10 μm (E) Force-Extension plot for oocyte spindle. The black line (representing the stretching process) and the yellow line (representing the retraction process) almost overlap with each other, which indicates that oocyte spindle displays elastic properties, and the stretching process is reversible (F) The doubling force of MI and MII oocytes shows similarity.

We assessed spindle stiffness following PBS exposure and found no significant difference between control spindles and those incubated in PBS for 30 minutes (Fig. S3 B). After one hour, a detectable decrease in spindle stiffness was observed (Fig. S3 B, an approximately 50% decrease). We conclude that the spindle is stable in PBS for 30 mins (the typical time of our mechanics experiments), but over additional time, the loss of certain components affects spindle stiffness. PBS is able to maintain spindle morphology for at least 30 minutes, enabling us to conduct further research on spindle mechanics *in vitro*.

A comparative analysis of spring constants – representing the force needed per length change, with units of force per length - revealed no significant difference between MI and MII oocyte spindles (MI spindle: 380 ± 35 pN/μm versus MII spindle: 450 ± 60 pN/μm) (Fig. S6). In addition, the length-doubling force, calculated by multiplying the spring constant by the spindle length to mitigate the influence of variable spindle length, exhibited no significant difference between the MI and MII spindles (MI spindle: 11000 ± 1000 pN versus MII spindle: 11000 ± 1600 pN) (Fig. 6 F). These results suggest a similarity in the organization and components of MI and MII spindles.

### Mechanics during oocyte spindle migration

Upon establishing the fundamental elastic properties of the oocyte spindles, we used the micropipette-micromanipulation method to investigate mechanics relevant to spindle migration during meiotic metaphase I. Under phase contrast, the spindle, observed as barrel-shaped with chromosomes in the middle, exhibited consistent features both *in vivo* and after isolation (Fig. 7 A). Since the migrating spindle is elastic, the spindle length changes before and after isolation can confirm pulling, but not pushing, forces are exerted onto the spindle as shown in Figure 1 D. Our results showed that the spindle length was significantly longer during spindle migration *in vivo* than after isolation (pre-isolation = 37.0 ± 0.6 μm; post-isolation = 33.9 ± 0.7 μm, *P* value <0.0001, paired test, Fig. 7 B left), suggesting that spindle migration is governed by a pulling force. Consistent with the pulling force model, the spindle width decreased inside cells (width before isolation = 16.2 ± 0.6 μm versus width after isolation = 20.1 ± 0.6 μm *P* value <0.0001, paired test) (Fig. 7 B right). This length increase and width decrease *in vivo* mirrors the stretching behavior for isolated spindles (Fig. 6 B, C and 7 F), further supporting the conclusion that the spindle is stretched within the oocyte.

**Figure 7.**
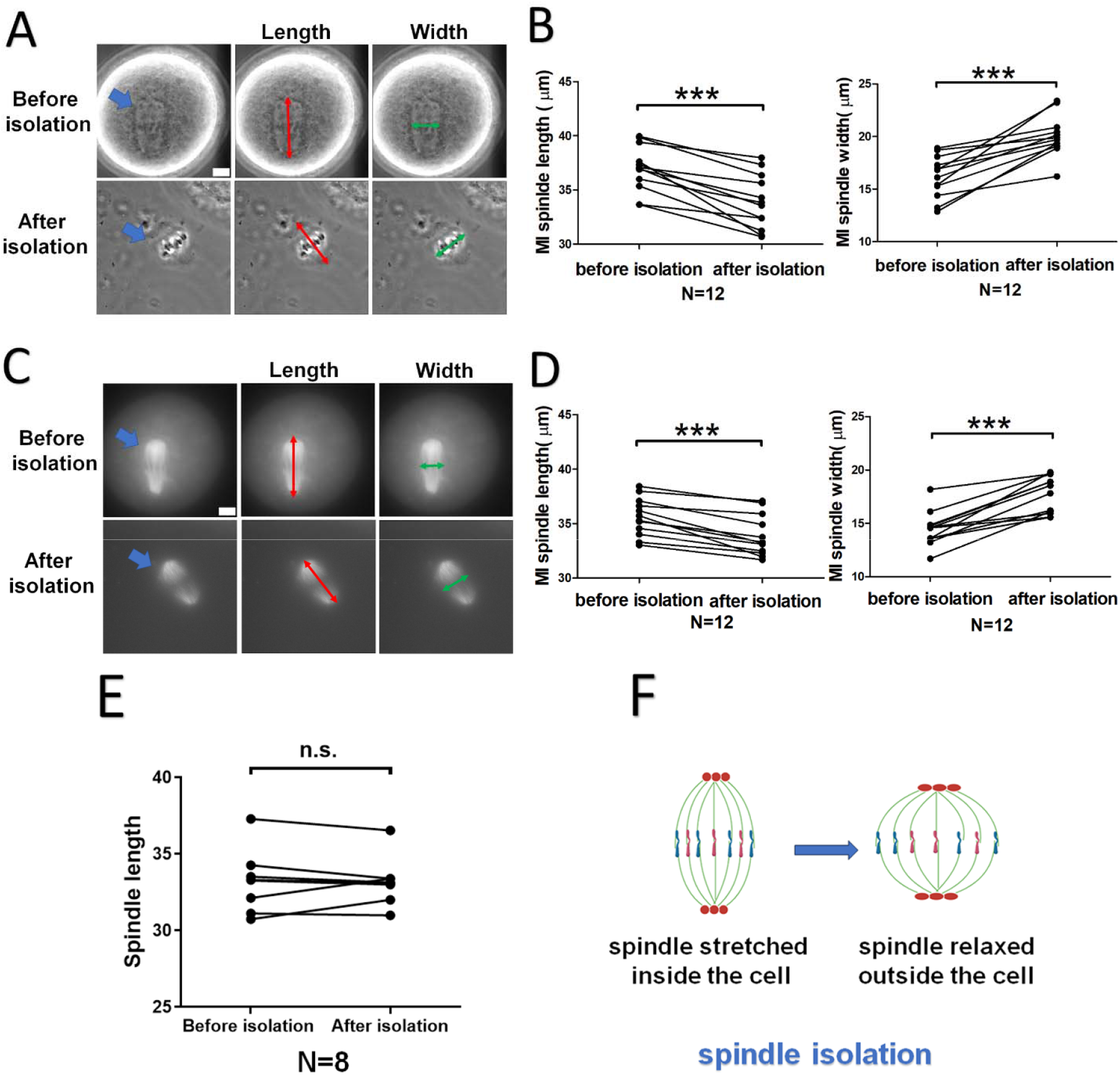
Spindle migration is governed by pulling forces rather than pushing forces. (A) A spindle before and after isolation under phase contrast. The spindle is visible both *in vivo* and post-isolation. Length and width were measured by connecting the midpoint of the ends or sides. Scale bar=10 μm (B) Spindle length and width changes before and after isolation under phase contrast. After isolation, spindle length decreases, and width increases, which indicates that the spindle is stretched inside the oocyte. (C) Spindle before and after isolation with microtubule staining. Scale bar=10 μm (D) Length and width changes of the immunostained spindle before and after isolation. Consistent with phase contrast observations, spindle length decreases, and width increases after isolation. (E) Spindle length after two minutes of cytochalasin D treatment does not change following isolation. (F) The MI spindle withstands stretching forces inside oocytes, and its migration is driven by pulling forces. After isolation, spindle length decreases while its width increases.

Microtubule staining validated the results (Fig. 7 C): upon isolation, the spindle length decreased (pre-isolation: 35.6 ± 0.5 μm versus post-isolation: 33.9 ± 0.5 μm, *P* value <0.0001, paired test), while the width increased (pre-isolation: 14.5 ± 0.5 μm versus post-isolation: 17.7 ± 0.5 μm, *P* value <0.0001, paired test) (Fig. 7 D). By subtracting the spindle length after isolation from the length before isolation, we obtained an average decrease in spindle length of 1.7 ± 0.3 μm. Measurement using the minimum bounding rectangle produced similar results (Fig. S7 A and B).

To assess the role of F-actin in spindle stretching, we compared spindle lengths before and after isolation in the presence of cytochalasin D. The results showed no significant difference between pre- and post-isolation spindle lengths, suggesting that F-actin contributes to spindle stretching within the cell (before isolation: 33.2 ± 0.7 μm versus after isolation: 33.2 ± 0.6, *P* = 0.9812, paired test) (Fig. 7E).

We conclude that spindle migration during meiosis is governed by pulling forces that create stretching and tension along the spindle axis. Based on the spindle spring constant and length change upon isolation, we estimated that the tension in the spindle during its migration from the oocyte center to the cortex region is 680 ± 120 pN (See Materials and methods). Given that spindle migration is a slow process with a speed of less than 0.1 μm/min (15), we assume that the forces at both ends of the spindle are balanced. Consequently, the active pulling force exerted by F-actin on spindle is also estimated to be approximately 680 pN.

## Discussion

Using a novel micromanipulation approach, we were able to isolate fresh intact spindles from mammalian oocytes at different stages, a process that proved to be uniquely feasible in oocytes while not possible in somatic cells. Surprisingly, the isolated spindles are stable in PBS, maintaining their structure for hour-long timescales.

This isolation method could provide a new model for spindle studies, especially for the MI spindle. Unlike bulk *Xenopus* egg extract-based methods, our method yields authentic spindles, maintains their original characteristics, allows observation under phase contrast, and requires fewer experimental treatments. Importantly, it allows for the examination of spindles at different substages aligned with the oocyte maturation timeline, presenting a valuable model for future spindle and chromosome investigations.

Spindle migration, a complex process influenced by numerous factors, such as microtubules, actin filaments, and motor proteins, has been considered using two hypotheses. The first posits a pulling force based on the fact that F-actin is enriched in the cortex region, and that myosin pulling on F-actin initiates spindle migration (11, 12). In contrast, a second pushing force hypothesis implicates a dynamic cytoplasmic F-actin meshwork nucleated by FMN2, which exerts the force necessary to relocate the spindle from the center to the cortex region (14–16). However, both hypotheses have been proposed based on observation, without substantive supporting evidence regarding force or mechanical measurements that could discern between them.

Our measurement of the spindle stiffness and length increase during migration provides evidence supporting and quantifying the pulling force hypothesis, i.e. that forces applied to the leading side of the spindle exceed those applied to the trailing side. Specifically, we determined that the oocyte spindle undergoes stretching forces inside cells, supporting the notion of a combination of F-actin/myosin network forces and metaphase cytoplasmic MTOC stretching the spindle (13, 35). Moreover, our calculated tension of approximately 680 pN within the spindle reinforces this. This force should be equal to the active pulling force exerted by F-actin if the speed of the spindle migration is constant. Given the potential subdivision of spindle migration into multiple substages (14, 36), whether this force is consistent throughout the entire process needs further investigation. In addition, it would be informative to determine whether spindle stretching is involved in regulating chromosome alignment and separation. Studying the relationship between spindle stretching and the force exerted by microtubules on chromosomes, which is crucial for proper chromosome separation, is an exciting area for further research.

One concern regarding our extraction experiments is that the spindles may have undergone alterations in their components after isolation, since PBS differs from the cytoplasmic environment. Isolating the spindle from the oocyte without losing any components appears to be unlikely, simply due to the complexity of the cytoplasmic composition. Such changes in spindle composition could lead to variations in spindle stiffness upon isolation. It is possible that a crucial structural component of the spindle is immediately lost during isolation, leading to a significant difference in spindle stiffness between vivo and vitro conditions. Transmitted light microscopy and stiffness measurements suggest that the spindles remain stable for at least 30 minutes, but not beyond 1 hour. This indicates that the molecular composition of the spindles may change post-isolation, most likely due to dilution of component concentrations outside the cell), leading to loss of some components through diffusion. These changes might only become apparent through transmitted light microscopy and overall structural alterations after some delay. However, it is also difficult to determine what components and what fraction of them are lost. There is currently no method to directly measure the stiffness of the spindle *in vivo* to test whether spindle stiffness remains constant. This would require directly assessing the force between the spindle and the cortex, which is challenging due to the difficulties of inserting equipment into oocytes and conducting such measurements. Alternative *in vivo* methods such as tension-dependent fluorescent proteins are a possibility for the future but remain difficult to calibrate and to measure anything expect one opening force. At present our experiment offers unique insights into the spindle mechanics during migration, offering an indirect way to estimate the force inside the oocytes, although the accurate force might be different due to loss of spindle components during isolation.

In summary, the isolation of fresh mammalian spindles from oocytes reported here is useful for study of spindle mechanics, positioning, and migration. Our method, when combined with genetic tools (e.g., siRNA, degron-based knockdowns, gene knockout or overexpression) and other pathologies like aging, offers potential to explore the functions of various microtubule-associated proteins (MAPs), shedding light on their roles in spindle positioning, migration, and orientation. The advantages of meiotic spindle isolation include the long duration of mammalian meiosis, providing a wider time window for precise substage studies. Additionally, the large size of oocytes facilitates microinjection for genetic or chemical manipulation.

## Materials and methods

### O-Ring culture well preparation

Prior to commencing the spindle experiments, it was imperative to prepare the O-ring culture wells. This involved rinsing a #1 coverslip with Sparkle (A.J. Funk & Co.) and gently wiping it with a Kimwipe (Kimberly-Clark). Subsequently, the coverslip was sprayed with ethanol and wiped clean with a Kimwipe, after which it was placed on a hotplate until thoroughly dry. Silicone O-rings (McMaster-Carr) were then dipped in molten paraffin wax and carefully positioned on the coverslip to create an O-Ring culture well. The resulting prepared O-Ring culture well underwent UV sterilization in a sterile hood for 40 minutes. All spindle experiments were conducted within the sterilized O-Ring culture well.

### Mouse oocyte collection and *in vitro* culture

CD1 mice (Charles River Laboratories) were used for this study. All mice were bred in the Pancoe Hall animal facility at Northwestern University. Mice were housed in a room under 12h dark and 12h light cycles. Animals have been fed on an ad libitum basis with free access to food and water. All protocols were approved by Northwestern University Institutional Animal Care and Use Committee.

Four-week-old female mice were selected for this study. Mice were euthanized by CO2 asphyxiation and dissected immediately to obtain ovaries. Dissected ovaries were rinsed several times in M2 medium (Sigma-Aldrich) containing 100 μM IBMX (Sigma-Aldrich). Following this, the ovaries were punctured using needles to release cumulus-oocyte complexes (COCs). The cumulus cells of the collected COCs were removed by pipetting several times to denude the GV-stage oocytes. To synchronize the oocytes at GV stage, the collected oocytes were soaked in M2 medium with 100 μM IBMX throughout the process. These denuded oocytes were then rinsed in M16 medium (Sigma-Aldrich) and finally transferred into a small drop of fresh M16 medium with or without chemical treatment. The culture medium drop was embedded in mineral oil in a culture dish (Fisher). The culture dish was put in a 5% CO2 incubator at 37⍰°C. Oocytes were cultured for different hours to reach different stages.

### Cell culture

Hela cells, MEF cells, and U2OS cells were cultured in DMEM complete medium composed of DMEM (Corning), 1% 100x penicillin/streptomycin (Corning), and 10% FBS (Hyclone). All the cells were cultured in a 37⍰°C incubator supplied with 5% CO_2_. Hela cells and MEF cells were passaged every 3-5 days. U2OS cells with GFP-labeled γ-tubulin cells were passaged every 7-9 days. All the cells were cultured for no more than 20 generation passages when they were used for experiments. Prior to the experiment, somatic cells were sub-cultured into O-Ring culture wells and cultured for 1-3 days under the condition of 37⍰°C and 5% CO_2_ to facilitate cell adhesion to the surface of O-Ring culture cell.

The TVI cell line was derived from newts (37). TVI cells were cultured in L-15 complete medium consisting of 50% L-15 medium, 39% sterile purified H_2_O, 10% FBS, and 1% 100x penicillin/streptomycin. The TVI cells were maintained at room temperature and underwent passage every two weeks. Regular weekly maintenance included the replacement of the culture medium.

### Spindle Micromanipulation

#### Micropipette preparation

Thin-wall glass capillaries were used to make micropipettes by mechanical pipette puller machine (Sutter P-97). Glass capillaries (6-inch length, OD 1.0 mm, with filament (TW100F-6)) were used to make capture pipettes (program 30: P=500; H=561; pull = 220; vel =200; time = 20). Glass capillaries (6-inch length, OD 1.0 mm, without filament (WPI TW100-6)) were used to make Triton and reagent spray pipettes (program 27: P=500; H=564; pull = 110; vel =110; time = 100). The forged pipette tips were subsequently cut to achieve different opening sizes: approximately 2 μm, 4 μm, and 50 μm for capture pipettes, Triton pipettes, and spray pipettes, respectively.

#### Micropipette treatment for spindle attachment

Glass capillaries (6-inch length, OD 1.0 mm, with filament) were treated with PC 2000 Plasma Cleaner (South Bay Technology) for 10 min to increase surface hydroxyls. Then glass capillaries were soaked in 1% of APTES (3-Aminopropyl) triethoxysilane) in methanol and rotated on a shaker overnight. Next, the APTES-coated capillaries were rinsed with methanol to wash out excess APTES. Following washing, the APTES-coated capillaries were dried in a chemical hood and used to make capture pipettes. The tips of capture pipettes had been soaked in 1% of glutaraldehyde for 1 h and then in methanol for 10 min to wash. Finally, the capture pipette tips were soaked in PBS for at least 10 min before utilization in experiments.

#### Cell lysis and spindle isolation

Spindle isolation was performed under an inverted microscope (IX-70; Olympus). Before the experiment, Triton pipettes were loaded with 0.05% Triton X-100 in PBS solution, capture pipettes were filled with PBS solution, and spray pipettes were filled with certain chemicals or antibodies. Cell lysis of somatic cells (Hela cells, MEF cells, TVI cells, and U2OS cells): the O-Ring culture well was placed on the microscope stage to search for a suitable cell. Once the target cell was identified, the triton pipette was moved close to the cell and then used to gently spray 0.05% Triton X-100 onto the cell’s surface.

Spindle isolation from oocytes was carried out by first treating post-culture oocytes in Acidic Tyrode’s solution for about 30 seconds to remove the zona pellucida. Oocytes were then transferred into an O-ring culture well containing 2ml of PBS for spindle isolation. For the cytochalasin D experiment, cytochalasin D was added to the PBS and the spindle length was measured under phase contrast before and after 2 min incubation. For isolation, oocytes and pipettes were positioned to the center of the field under the 10X objective lens. A higher magnification objective lens (60X with a 1.5× magnification pullout) was then used to obtain a closer view. Next, 0.05% Triton X-100 was sprayed onto the surface of oocytes treated with Acidic Tyrode’s solution. The Triton X-100 treatment broke the oocyte membrane, allowing the spindle to flow out of the cells spontaneously. If spindle did not flow out, it can was squeezed out using a pipette to gently put pressure onto the oocyte. The capture pipette was used to grab one pole of the spindle and drag it into an empty spot within the well. Another capture pipette was used to capture the other pole of the spindle. In this way, the two poles of spindle were grabbed and held by two capture pipettes, making it ready for stiffness measurement.

#### Spindle stiffness measurement

After capturing the spindle poles with two capture pipettes, the angle of the pipettes holding the spindle was adjusted to ensure that the spindle axis was perpendicular to the pipette. The pipette was fixed on a motorized manipulator and controller (MP-285, Sutter Instruments). A LabView (National Instruments) program was utilized to control the movement of the pipette. The pipette was moved perpendicularly to metaphase plate at a constant rate of 0.20⍰μm/s with 0.04⍰μm steps for 30s to stretch the spindle. The actual movement and location change of the two pipettes were tracked and recorded under the microscope using a CCD camera and frame grabber. Each stretching process was repeated up to 6 times. Entire measurement for one spindle was done within 1 hour to maintain the spindle’s initial organization. After the measurement, spring constant of the spindle was obtained by multiplying the spring constant of the pipette by the ratio of the pipette deflection to the spindle deformation determined from the initial linear mechanical response (K_spindle_=K_pipette_* D). Doubling force was obtained by multiplying spindle spring constant by spindle length (F_double_= L_0_*K_pipette_). K_spindle_ was the spindle spring constant obtained by calculation, F_double_ was the doubling force obtained by calculation, D was the ratio of the force-measurement deflection to spindle stretching, K_pipette_ was the pipette spring constant obtained by pipette calibration, L_0_ was spindle length obtained by microscope.

#### Calculation of force for spindle migration

The change in spindle length, L_change_ is calculated by measuring the difference in spindle length before isolation and after isolation. The force driving spindle migration is obtained via multiplying L_change_ by the spindle spring constant, K_spindle_. The standard error is chosen by selecting the higher percentage between L_change_ and K_spindle_. Therefore, the force is computed by multiplying 1.7 ± 0.3 μm and approximately 400 pN/μm (380 pN/μm). Consequently, the estimated force is 680 ± 120 pN.

#### Micropipette calibration

A suitable calibrated pipette was obtained by a series of stepwise calibratinos off gradually less stiff pipettes. A relatively stiff master pipette (roughly 1000 pN/micron spring constant) served as the initial reference and underwent calibration using a nanonewton-sensitivity electronic force transducer (FemtoTools). A first calibrated pipette with a lower spring constant was calibrated using the master pipette. A second calibrated pipette with a yet lower spring constant was calibrated using the first calibrated pipette. This second calibrated pipette had a spring constant in the 100 pN/micron range and was suitable for calibrating similar-stiffness experimental force-measuring pipettes.

#### Microspraying of chemicals and antibodies

The spray pipette was loaded with 10 μl of the respective antibodies or chemicals using a pump, ensuring the careful exclusion of bubbles. Next, the spray pipette was brought close to the spindle under a 10X non-contact objective lens. The antibody and chemicals in the spray pipette were then sprayed onto the spindle gently at a constant rate. The whole spray process takes 10 min unless specifically mentioned. For microtubule staining, 2 μl of anti-α-tubulin-FITC (Sigma-Aldrich) diluted in 48 μl of PBS buffer were applied. For Hoechst staining, stock solution was made by diluting 1 μl of Hoechst (33342) in 10 ml PBS. Subsequently, 5 μl of stock solution were added to PBS with other chemicals and antibodies to reach a total of 50 μl spray solution if DNA staining was needed. After spraying, the spindle was relocated away from the sprayed region to prevent the remnant signal of antibodies. This precaution was taken to mitigate background noise caused by the unbound antibody around the spindle or antibodies attached to the culture cell surface.

#### Live-cell microtubule staining

ViaFluor 488 Live Cell Microtubules staining Kit (Biotium) was used to stain microtubules in live oocytes and somatic cells. For oocytes, 0.2 μl of ViaFluor Live Cell Microtubule Stain and 0.4 μl of Verapamil HCl were added to 400 μl prewarmed M16 Medium. The post-culture oocytes were treated in the M16 media for 3 min at 37⍰°C in a 5% CO_2_ incubator. For somatic cells, 1 μl of ViaFluor Live Cell Microtubule Stain and 2 μl of Verapamil HCl were added to 10 ml of DMEM complete medium to make a microtubule staining medium. Staining medium was prewarmed for 30 min at 37⍰°C. Somatic cells were treated with the microtubule staining medium for 3 min at 37⍰°C in a 5% CO_2_ incubator.

#### Laser ablation of mcMTOCs

Preparation and oocyte culture: Cep192-eGFP reporter (13) were generated and maintained in experimental guide accordance with the University of Missouri. All mice were kept in a photoperiod of 12h light/dark cycle at 21°C, 55% humidity, food and water ad libitum. Full-grown GV oocytes were isolated from the Cep92-eGFP female ovaries (6-8 weeks old) and incubated in Chatot, Ziomek, and Bavister (CZB) medium (38) supplemented with SiR-tubulin (Cytoskeleton #NC0958386) at 37°C with 5% CO2.

mcMTOCs depletion conducted by two-photon laser ablation: Using a Leica TCP SP8 two-photon inverted microscope, equipped with a microenvironmental chamber to control the temperature and CO2, the mcMTOCs were ablated using an 820nm wavelength laser according to published papers (13, 35). The mcMTOCs were marked by small squares and then exposed to a laser beam. The focal plane was moved allowing us to deplete all mcMTOCs present in the oocyte. Control group was exposed to the same conditions except random areas in the cytoplasm of similar size to mcMTOCs, adjacent but not overlapping with mcMTOCs, were ablated. The oocytes were imaged live following laser ablation of mcMTOCs. The length of metaphase I spindle was measured using NIH image J software (National Institute of Health, Bethesda, MD, USA)

#### Statistical analysis

All the analysis was done in GraphPad Prism. Data of means and standard errors were obtained. Unpaired two-tailed t-tests or paired sample t-tests were used for statsistical significance depending on the sample types. Comparisons between different groups of samples were considered as significant when *P < 0.05, **P < 0.01 and ***P < 0.001.

## Data availability

Data for this study is available in the published article itself and its supplementary materials.

## Acknowledgments

We would like to thank Dr. Caroline Mauvezin for providing the U2OS cell line. We want to thank Dr. Karen Schindler and all the members of the Qiao and Marko groups, especially Dr. Ronald J. Biggs for their technical support and comments on this manuscript.

This work was supported by National Institutions of Health (NIH)R00 HD082375 and (NIH)R01 GM135549, (NIH)UM1-HG011536, (NIH)R01-GM105847, U54 CA193419, and U54 CA203000.

## Author contributions

N. Liu, H. Qiao and J. F. Marko conceived the project; N. Liu designed the experiments; N. Liu analyzed the data and performed the experiments; A. Balboula, R. Kawamura and W. Qiang provided technical assistance in the experiments and manuscript editing; and N. Liu, H. Qiao and J. F. Marko wrote the manuscript. All authors read and approved of the final manuscript.

## References

1. S. Dumont, T. J. Mitchison, Force and Length in the Mitotic Spindle. Curr. Biol. 19, R749–R761 (2009).

2. M. J. Lohka, Y. Masui, Formation in vitro of sperm pronuclei and mitotic chromosomes induced by amphibian ooplasmic components. Science (80-.). 220, 719–721 (1983).

3. F. Verde, J.C. Labbé, M. Dorée, E. Karsenti, Regulation of microtubule dynamics by cdc2 protein kinase in cell-free extracts of Xenopus eggs. Nature 343, 233–238 (1990).

4. M. I. Lohka, J. L. Maller, Induction of nuclear envelope breakdown, chromosome condensation, and spindle formation in cell-free extracts. J. Cell Biol. 101, 518–523 (1985).

5. A. Desai, A. Murray, T. J. Mitchison, C. E. Walczak, Chapter 20 The Use of Xenopus Egg Extracts to Study Mitotic Spindle Assembly and Function in Vitro. Methods Cell Biol. 61, 385–412 (1998).

6. R. Heald, et al., Self-organization of microtubules into bipolar spindles around artificial chromosomes in Xenopus egg extracts. Nature 382, 420–425 (1996).

7. R. Gibeaux, et al., Paternal chromosome loss and metabolic crisis contribute to hybrid inviability in Xenopus. Nature 553, 337–341 (2018).

8. Y. Shimamoto, T. M. Kapoor, Microneedle-based analysis of the micromechanics of the metaphase spindle assembled in xenopus laevisegg extracts. Nat. Protoc. 7, 959–969 (2012).

9. T. Fukuyama, et al., Morphological growth dynamics, mechanical stability, and active microtubule mechanics underlying spindle self-organization. Proc. Natl. Acad. Sci. U. S. A. 119, 1–11 (2022).

10. E. D. Salmon, S. M. Wolniak, Taxol stabilization of mitotic spindle microtubules: Analysis using calcium induced depolymerization. Cell Motil. 4, 155–167 (1984).

11. D. H. Park, L. S. Rose, Dynamic localization of LIN-5 and GPR-1/2 to cortical force generation domains during spindle positioning. Dev. Biol. 315, 42–54 (2008).

12. M. Schuh, J. Ellenberg, A New Model for Asymmetric Spindle Positioning in Mouse Oocytes. Curr. Biol. 18, 1986–1992 (2008).

13. D. Londoño-Vásquez, K. Rodriguez-Lukey, S. K. Behura, A. Z. Balboula, Microtubule organizing centers regulate spindle positioning in mouse oocytes. Dev. Cell 57, 197-211.e3 (2022).

14. K. Yi, et al., Sequential actin-based pushing forces drive meiosis I chromosome migration and symmetry breaking in oocytes. J. Cell Biol. 200, 567–576 (2013).

15. X. Duan, et al., Dynamic organelle distribution initiates actin-based spindle migration in mouse oocytes. Nat. Commun. 11, 1–15 (2020).

16. H. Li, F. Guo, B. Rubinstein, R. Li, Actin-driven chromosomal motility leads to symmetry breaking in mammalian meiotic oocytes. Nat. Cell Biol. 10, 1301–1308 (2008).

17. J. F. Casella, M. D. Flanagan, S. Lin, Cytochalasin D inhibits actin polymerization and induces depolymerization of actin filaments formed during platelet shape change. Nat. 1981 2935830 293, 302–305 (1981).

18. C. So, et al., A liquid-like spindle domain promotes acentrosomal spindle assembly in mammalian oocytes. Science (80-.). 364 (2019).

19. R. J. Biggs, N. Liu, Y. Peng, J. F. Marko, H. Qiao, Micromanipulation of prophase I chromosomes from mouse spermatocytes reveals high stiffness and gel-like chromatin organization. Commun. Biol. 3, 1–7 (2020).

20. N. Liu, W. Qiang, P. W. Jordan, J. F. Marko, H. Qiao, Cell cycle and age-related modulations of mouse chromosome stiffness. Elife 13 (2025).

21. P. T. Tran, P. Joshi, E. D. Salmon, How tubulin subunits are lost from the shortening ends of microtubules. J. Struct. Biol. 118, 107–118 (1997).

22. G. Halet, J. Carroll, Rac Activity Is Polarized and Regulates Meiotic Spindle Stability and Anchoring in Mammalian Oocytes. Dev. Cell 12, 309–317 (2007).

23. J. Brugués, D. Needleman, Physical basis of spindle self-organization. Proc. Natl. Acad. Sci. U. S. A. 111, 18496–18500 (2014).

24. K. E. Sawin, T. J. Mitchison, Poleward microtubule flux in mitotic spindles assembled in vitro. J. Cell Biol. 112, 941–954 (1991).

25. R. Kawamura, et al., Mitotic chromosomes are constrained by topoisomerase II-sensitive DNA entanglements. J. Cell Biol. 188, 653–663 (2010).

26. A. R. Strom, et al., HP1α is a chromatin crosslinker that controls nuclear and mitotic chromosome mechanics. Elife 10, 1–30 (2021).

27. S. Busson, D. Dujardin, A. Moreau, J. Dompierre, J. R. De Mey, Dynein and dynactin are localized to astral microtubules and at cortical sites in mitotic epithelial cells. Curr. Biol. 8, 541–544 (1998).

28. J. Rosenblatt, L. P. Cramer, B. Baum, K. M. McGee, Myosin II-dependent cortical movement is required for centrosome separation and positioning during mitotic spindle assembly. Cell 117, 361–372 (2004).

29. M. Carminati, et al., Concomitant binding of Afadin to LGN and F-actin directs planar spindle orientation. Nat. Struct. Mol. Biol. 23, 155–163 (2016).

30. M. Okumura, T. Natsume, M. T. Kanemaki, T. Kiyomitsu, Dynein–dynactin–NuMA clusters generate cortical spindle-pulling forces as a multiarm ensemble. Elife 7, 1–24 (2018).

31. D. J. Sharp, G. C. Rogers, J. M. Scholey, Microtubule motors in mitosis. Nature 407, 41–47 (2000).

32. M. Schuh, J. Ellenberg, Self-Organization of MTOCs Replaces Centrosome Function during Acentrosomal Spindle Assembly in Live Mouse Oocytes. Cell 130, 484–498 (2007).

33. B. J. Schnackenberg, A. Khodjakov, C. L. Rieder, R. E. Palazzo, The disassembly and reassembly of functional centrosomes in vitro. Proc. Natl. Acad. Sci. U. S. A. 95, 9295–9300 (1998).

34. R. B. Nicklas, C. A. Koch, Chromosome micromanipulation. 3. Spindle fiber tension and the reorientation of mal-oriented chromosomes. J. Cell Biol. 43, 40–50 (1969).

35. D. Londoño-Vásquez, A. Jurkevich, A. Z. Balboula, Multi-Photon Laser Ablation of Cytoplasmic Microtubule Organizing Centers in Mouse Oocytes. J. Vis. Exp. 2022, 1–10 (2022).

36. M. Bezanilla, P. Wadsworth, Spindle Positioning: Actin Mediates Pushing and Pulling. Curr. Biol. 19, R168–R169 (2009).

37. J. F. Marko, M. Poirier, S. Eroglu, D. Chatenay, Reversible and irreversible unfolding of eukaryote chromosomes by force. Annu. Int. Conf. IEEE Eng. Med. Biol. -Proc. 1, 79 (1999).

38. C. L. Chatot, C. A. Ziomek, B. D. Bavister, J. L. Lewis, I. Torres, An improved culture medium supports development of random-bred 1-cell mouse embryos in vitro. Reproduction 86, 679–688 (1989).

