## Supplemental Figures for "Isolation and manipulation of meiotic spindles from mouse oocytes reveals migration regulated by pulling force during asymmetric division"

**Supplemental materials**

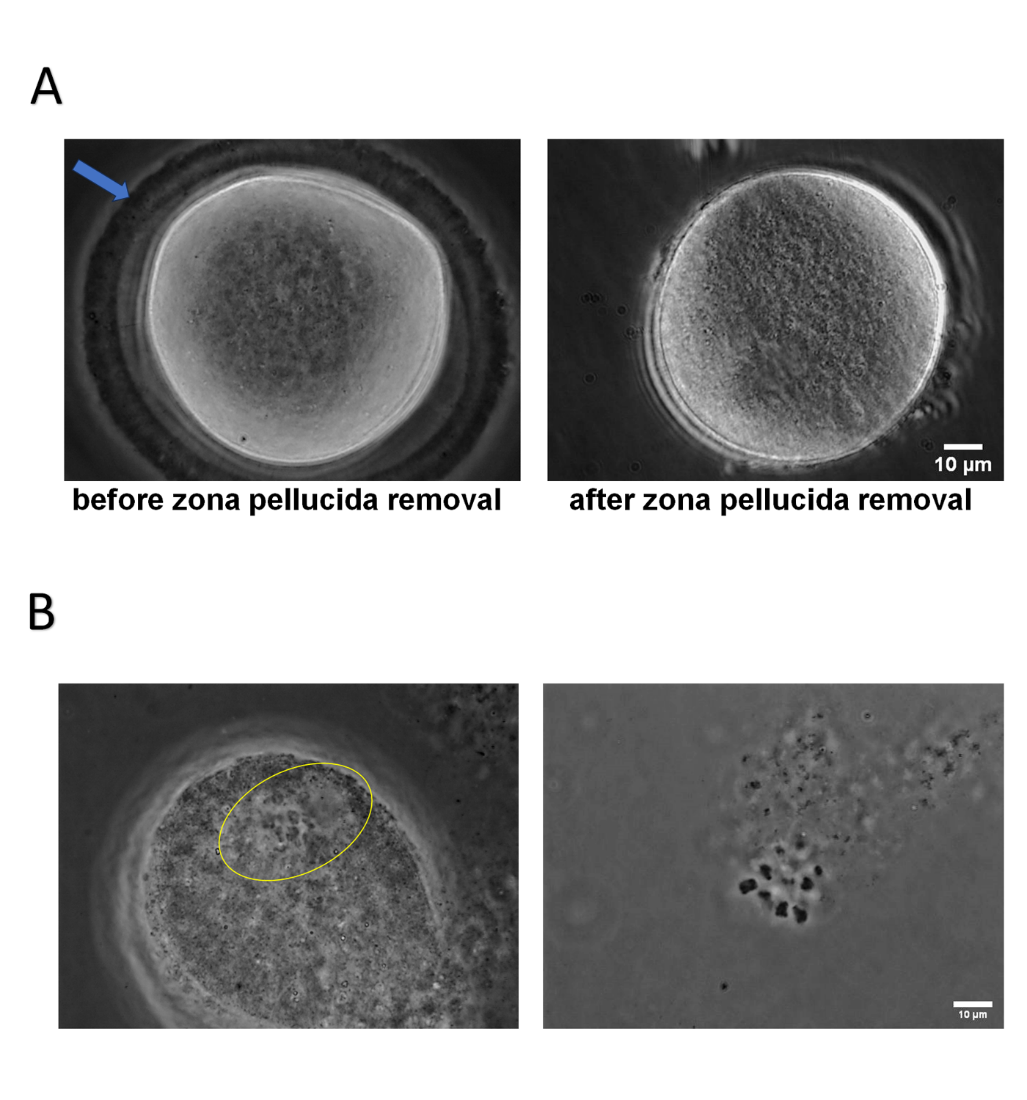

Figure S1. **Zona pellucida removal and spindle isolation.** (A) MI oocyte before and after zona pellucida removal. (B) Spindle isolation failure. Left panel: spindle inside oocyte (circled with yellow) unable to be isolated. Right panel: spindle formation delayed. Typical barrel-shaped spindle could not be seen.

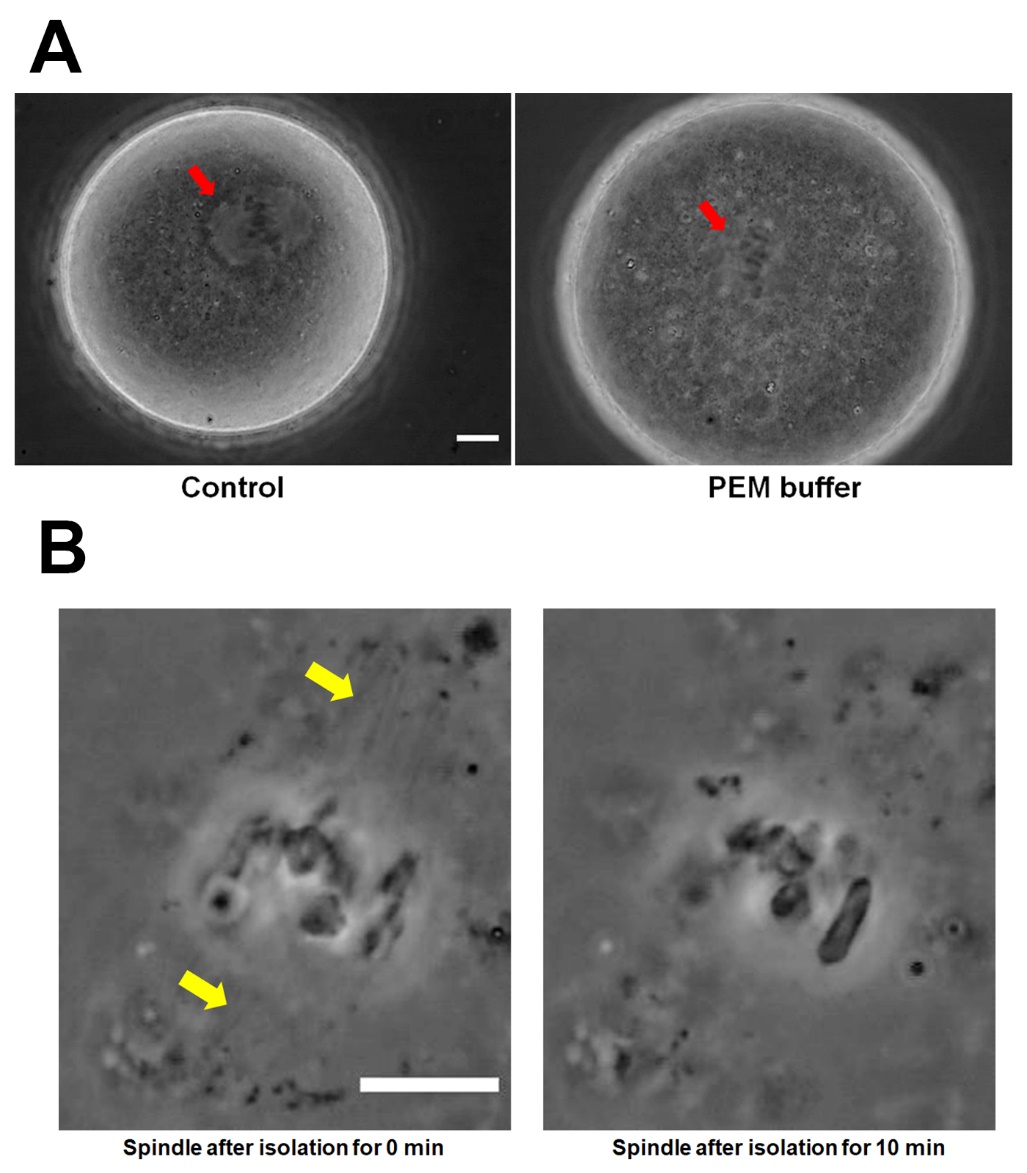

Figure S2. **PEM buffer is detrimental to oocytes and spindles.** (A) PEM buffer is toxic to oocytes. Oocytes in PEM buffer appear larger and darker compared to those in PBS control. Moreover, the spindle (red arrow) lacks the clear outline seen in PBS. Scale bar=10 μm (B) The spindle disassembles rapidly in osmotically balanced PEM solution (PEM+150mM NaCl). Right after isolation, the microtubule structure in the spindle is visible (yellow arrow), but becomes undetectable affter 10min. Scale bar=10 μm

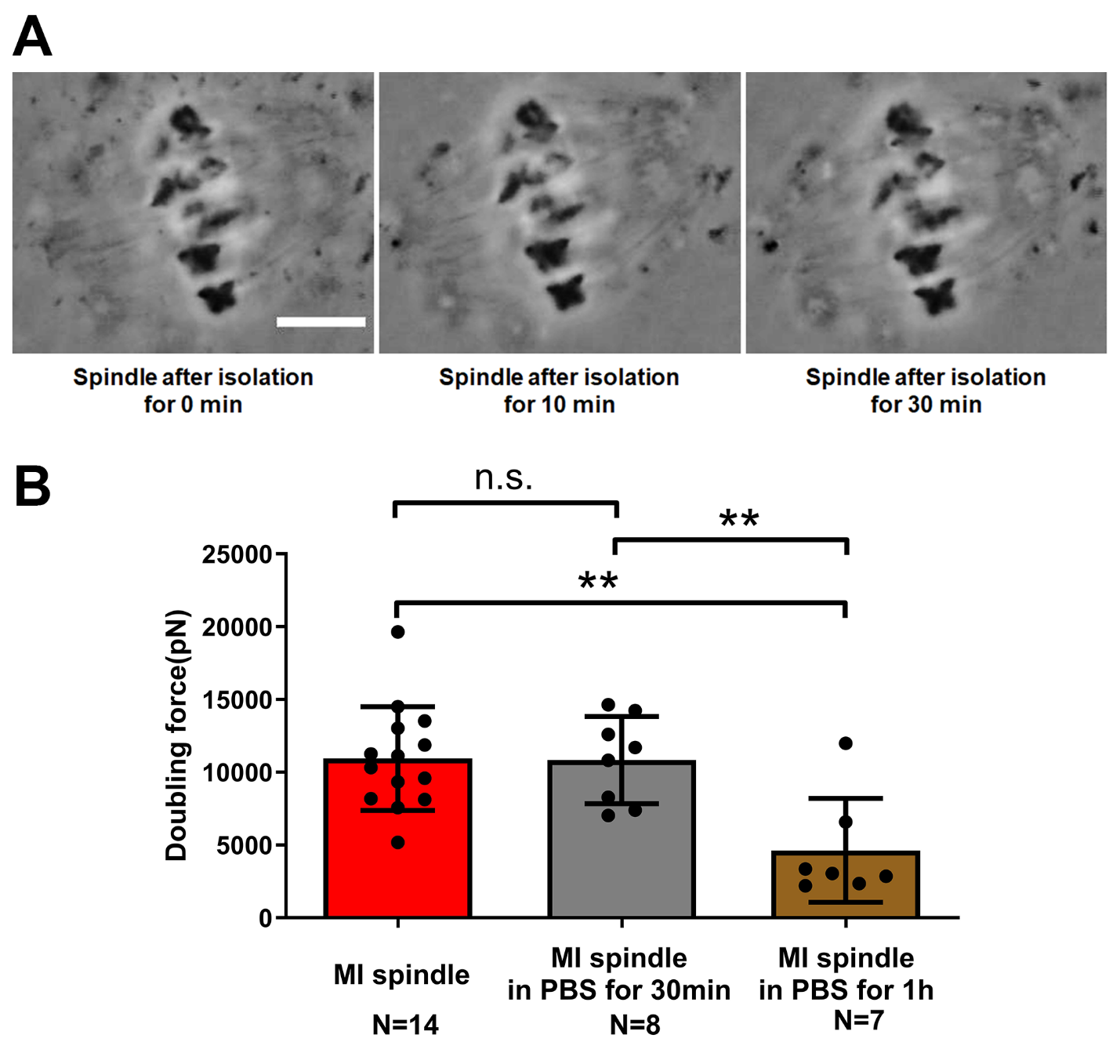

Figure S3. **Isolated spindles are stable in PBS solution.** (A) Isolated spindle can matin its morphology for at least 30 min in PBS solution. Scale bar=10 μm (B) Spindle stiffness changes over time after isolation from oocyte in PBS solution. The spindle stiffness of MI spindle for 0h (control) (11000 ± 1000 pN) is significantly higher than that of spindles in PBS for 1h (5000 ± 1000 pN) and shows no significant difference compared to MI spindles in PBS for 30 min (11000 ± 1000 pN). Additionally, the spindle stiffness of MI spindles in PBS for 30 minutes is higher than that of spindles in PBS for 1h

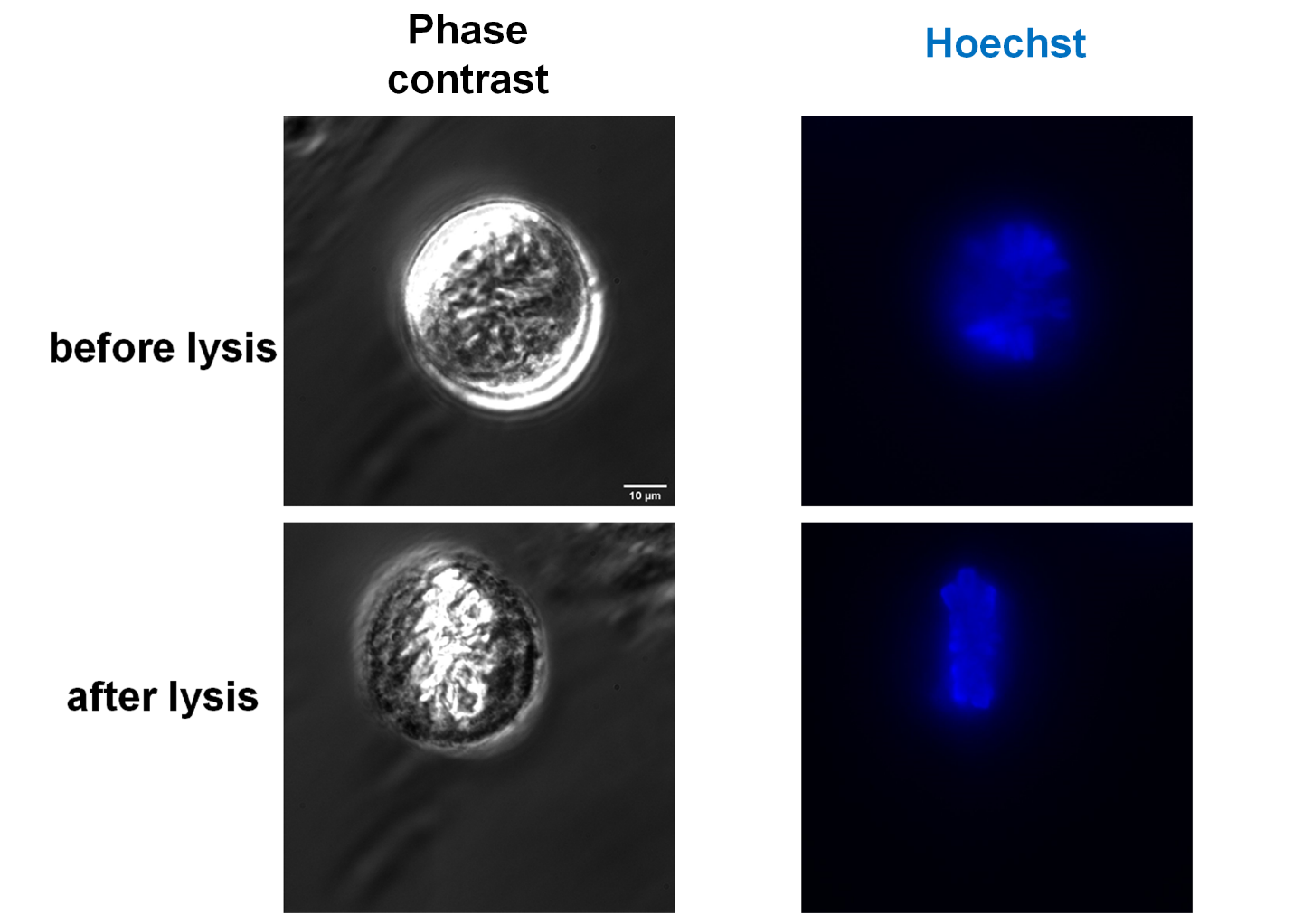

Figure S4. **TVI cell before and after cell lysis.** Left panel: phase contrast; Right panel: Hoechst staining of DNA/chromosomes. Chromosomes aligned linearly in the middle of the cell indicate that the cell is at metaphase. stage.

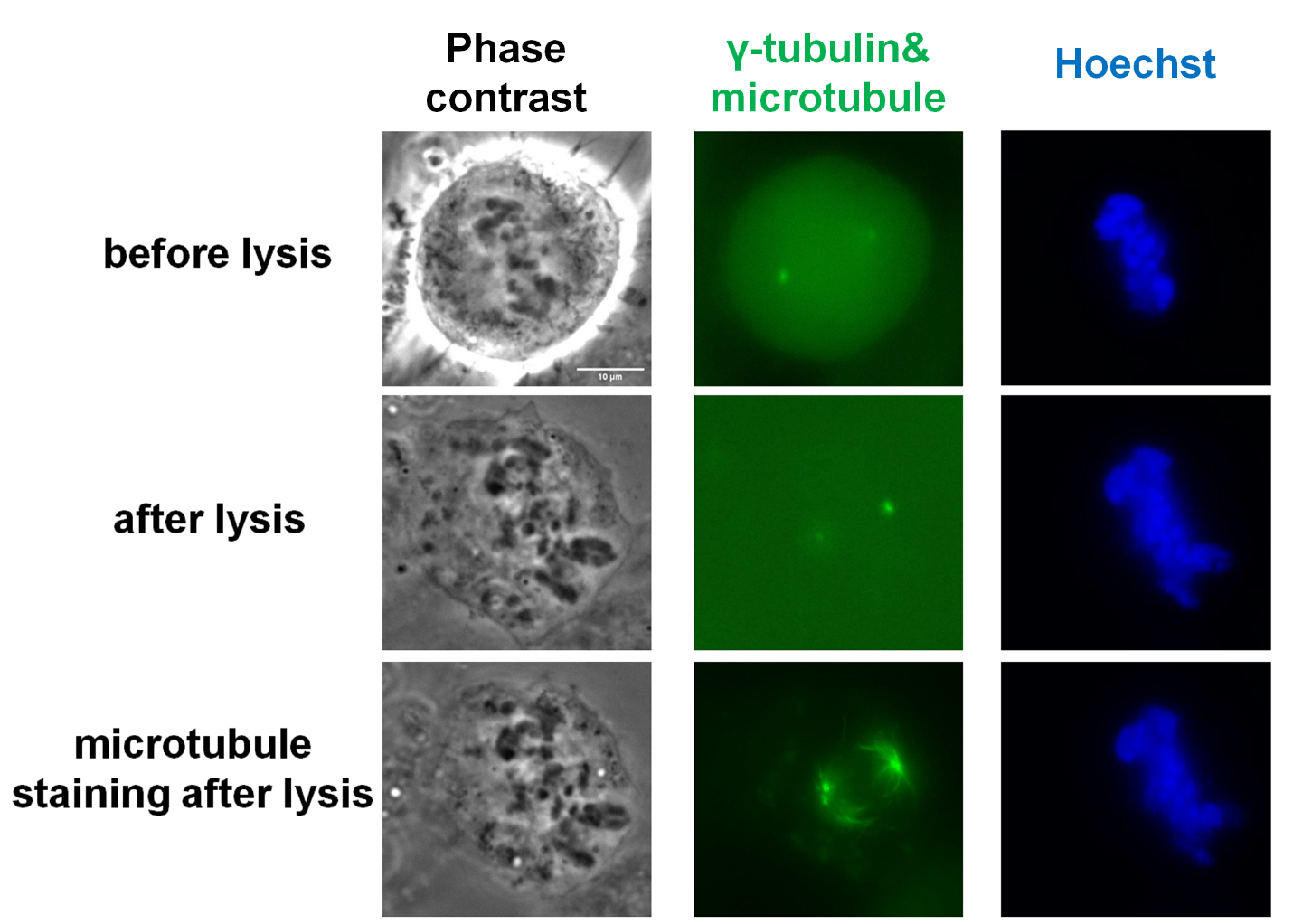

Figure S5. **Microtubule immunostaining of U2OS cell before and after cell lysis.** Left panel: phase contrast. Middle panel: γ-tubulin (GFP) & microtubule (if stained). Right panel: Hoechst. The spindle is stained after cell lysis instead of in vivo staining. In this way, we can rule out the possibility that the spindle should be disrupted after cell lysis without stabilization, but the microtubule dye could stabilize the spindle, thus maintaining the intact spindle structure after cell lysis.

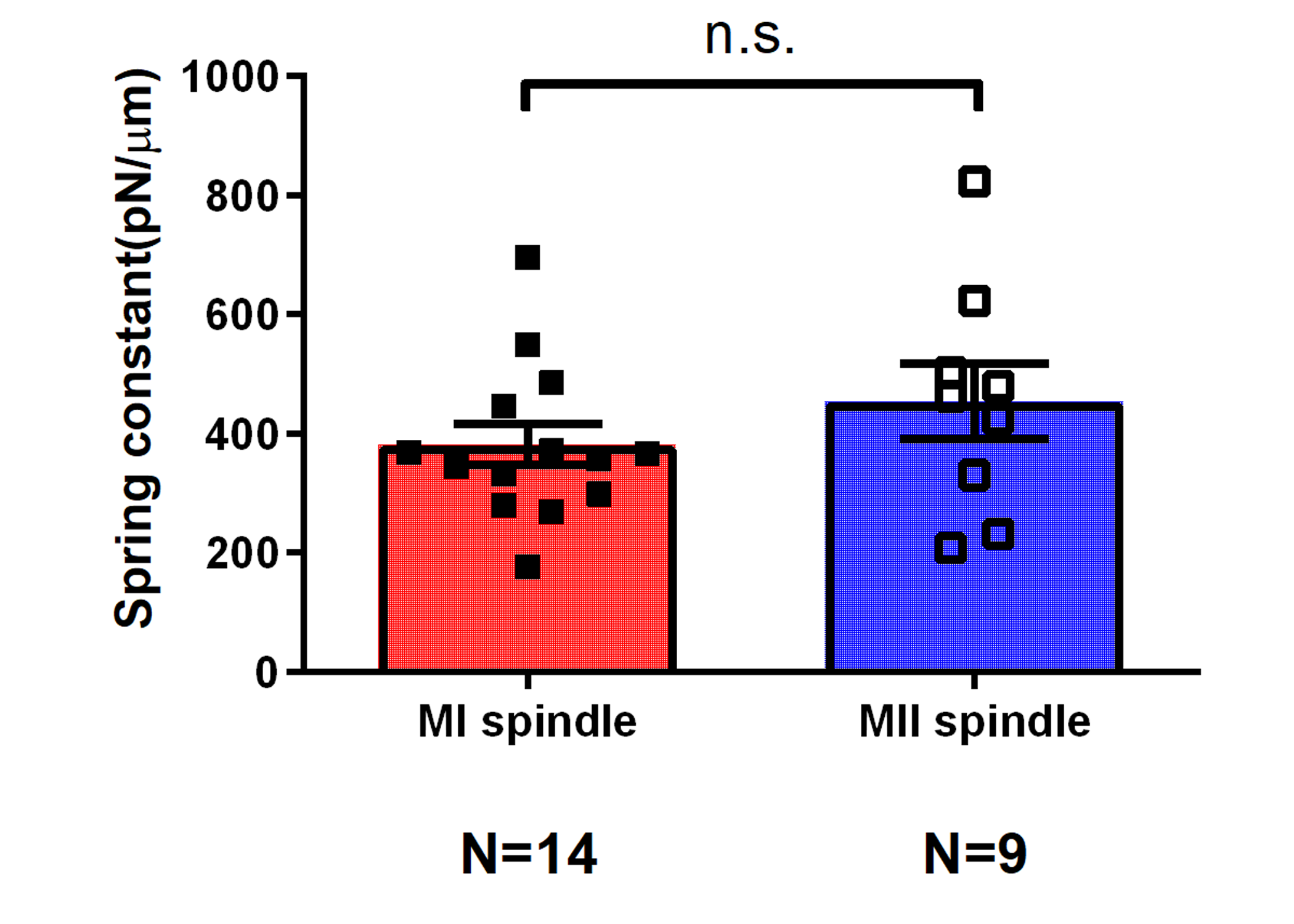

Figure S6. **Spindle spring constant measurement.** The spring constants of MI spindle and MII spindle is not statistically different.

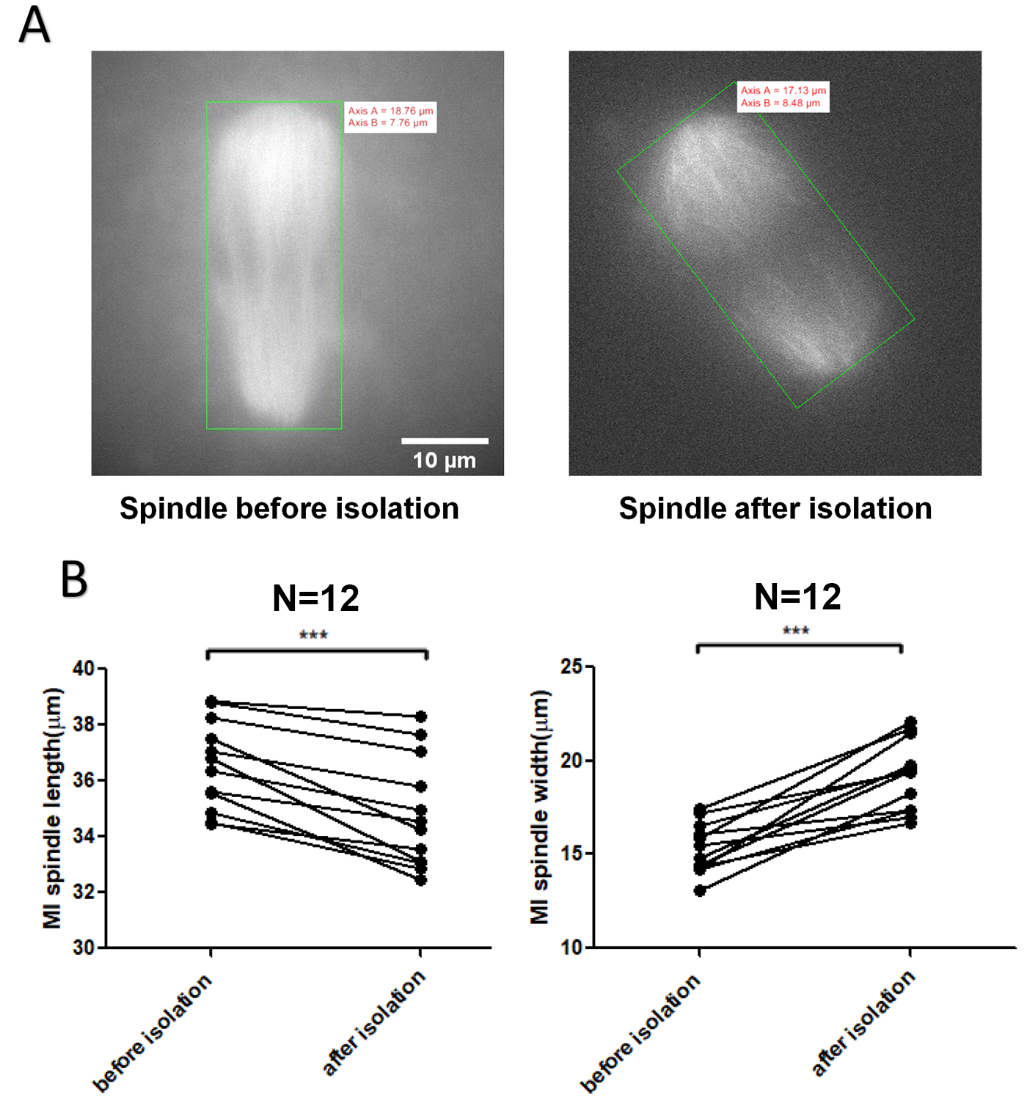

Figure S7. **Spindle length and width measurement by minimum bounding rectangle.** (A) MI spindle length and width measurement by drawing the smallest rectangle which could enclose the whole spindle. Axis A is a half of spindle length. Axis B is a half of spindle width. (B) MI spindle length and width change before and after isolation. After isolation, the spindle length is decreased while the spindle width is increased.

Video 1: **Movie monitoring the spindle morphology change over time after isolation from oocyte in PBS solution.**

Video 2: **Time-lapse movie of strong stretching of the spindle from MI oocyte under the phase contrast.** When the spindle undergoes stretching, its length increases while the width decreases, indicating that the spindle is elastic and has a deformable gel-like structure. The extension shown is larger than that used in actual runs used to measure elasticity.
